# ZetaSuite, A Computational Method for Analyzing Multi-dimensional High-throughput Data, Reveals Genes with Opposite Roles in Cancer Dependency

**DOI:** 10.1101/2021.08.29.458095

**Authors:** Yajing Hao, Changwei Shao, Guofeng Zhao, Xiang-Dong Fu

## Abstract

The rapid advance of high-throughput technologies has enabled the generation of two-dimensional or even multi-dimensional high-throughput data, e.g., genome-wide siRNA screen (1^st^ dimension) for multiple changes in gene expression (2^nd^ dimension) in many different cell types or tissues or under different experimental conditions (3^rd^ dimension). We show that the simple Z-based statistic and derivatives are no longer suitable for analyzing such data because of the accumulation of experimental noise and/or off-target effects. Here, we introduce ZetaSuite, a statistical package designed to score and rank hits from two-dimensional screens, construct regulatory networks based on response similarities, and eliminate off-targets. Applying this method to two large cancer dependency screen datasets, we identify not only genes critical for cell fitness, but also those required for constraining cell proliferation. Strikingly, most of those cancer constraining genes function in DNA replication/repair checkpoint, suggesting that cancer cells also need to protect their genomes for long-term survival.

## Main

Genome-wide screen by RNA interference (with siRNA or shRNA) ^1–3^ or CRISPR/Cas (with sgRNA) ^4–6^ has been extensively employed to identify global regulators in specific biological pathways. Screen strategies with either arrayed or pooled targeting RNAs have been developed, but most applications score a single functional readout, which we here refer to as one-dimensional (many-to-one) screen. The array-based strategy has been extended to examine multiple functional parameters in response to each treatment, known as multi-content (many-to-a few) screen ^1,7^. In the deep sequencing era, the array-based strategy has further evolved toward two-dimensional high throughput (many-to-many) screen because of the feasibility to perform functional perturbations and monitor functional consequences, both in a high throughput fashion, as exemplified by the HTS^2^ screen platform to monitor a gene signature comprising a set of specific genes, rather than a single gene ^8,9^. Pooled shRNA or sgRNA libraries have also been used to treat hundreds of cell lines to deduce cancer dependencies ^10–13^, representing another format of two-dimensional screen. It may soon become no longer cost-prohibitive to perform genome-wide functional perturbation (1^st^-) and measure genome-wide response (2^nd^-) in many different cell types or tissues (3^rd^-dimension).

The advance in high throughput technologies is frequently accompanied with the demand for developing new analytical tools to treat the data of increasing complexity. For one-dimensional large-scale screen, t-test, Z-statistic or Robust Z, or strictly standardized mean difference (SSMD) or Robust SSMD ^14^ have been common choices to identify screen hits, depending on the availability of replicates and build-in positive and/or negative controls [see ^15^ for choosing the most suitable method for processing data from different types of screen]. However, as demonstrated in the current study, these statistical approaches are not optimal for analyzing two-dimensional high throughput screen data due to the accumulation of experimental errors and off-target effects.

Off-target effects remain a major challenge in analyzing screen results with siRNA, shRNA or sgRNA. A general strategy to meet this challenge is to increase the number of targeting RNAs against each gene and aggregate enriched hits to reflect the collective effect, as with RSA ^16^, RIGER ^17^, and more recently, MAGeCK ^18^. Furthermore, ATARiS ^19^ and DEMETER2 ^20^ have been designed to remove targeting RNAs that likely cause off-target effects. In ATARiS, for example, a set of targeting RNAs is each tested on multiple samples to identify those that show the overall similarity across the samples, assuming that the rest likely cause off-target effects. In DEMETER ^11^ or DEMETER2 ^20^, targeting RNAs that may cause off-target effects are filtered by using their sequences in the corresponding seed region to calculate potential microRNA-like effects on off-targets. Since numerous microRNAs do not strictly follow such seed rule ^21,22^. it remains unclear to what extent this approach actually helps reduce the off-target effects. Importantly, as most of these approaches require a large number of targeting RNAs per gene (around 15 to 20), they are not applicable to screens with traditionally arrayed siRNAs, typically consisting of 4 to 6 targeting RNAs per gene. In fact, the increased sequence complexity in each set of targeting RNAs may also elevate the probability in off-targeting, thus causing more artifacts than eliminated in some cases.

In this study, we introduce a new ZetaSuite designed to address multiple challenges in two or multi-dimensional high throughput screens with either pooled or arrayed libraries, which is perceived to have increasing utilities in modern biological research due to the ever-increasing power of deep sequencing. ZetaSuite (available at https://github.com/YajingHao/ZetaSuite) can identify and rank hits at a full Z range, rather than on an arbitrarily chosen cutoff, to differentiate between positives and negatives. When a large number of true positive and negative controls is available, ZetaSuite can further draw a support vector machine (SVM) learning curve to maximally separate positives from negatives. We first develop ZetaSuite by using in-house two-dimensional siRNA screen data designed to identify global splicing regulators, demonstrating that the core spliceosome components are the most dominant class of regulators for alternative splicing in mammalian cells. We next apply ZetaSuite to the existing cancer dependency maps, revealing not only genes essential for cancer cells to survive, as reported earlier ^10^, but also genes that act to prevent excessive cancer cell proliferation, most corresponding to those involved in DNA replication/repair checkpoint control. These findings demonstrate the broad utility of ZetaSuite in processing multi-dimensional high throughput data to expose critical regulatory pathways.

## Results

### Overview of the ZetaSuite frame workflow

ZetaSuite is a statistical and computational framework initially developed to process the data from a siRNA screen for global splicing regulators [see ^23^]. In this screen, we interrogated ∼400 endogenous alternative splicing (AS) events by using an oligo ligation-based strategy to quantify their responses to 18,480 pools of siRNAs against annotated protein-coding genes in the human genome (Supplementary Fig. 1a). We performed deep sequencing on bar-coded and pooled samples from individually treated wells in 384-well plates to generate digital information on interrogated mRNA isoforms, and by comparing to internal non-specific siRNA treated negative controls, we were able to quantify induced exon inclusion or skipping for each AS event (similar to up- and down-regulated genes from a typical RNA-seq experiment). The resultant data matrix resembles those produced by high-content screens, parallel genome-wide screens, or any screens that monitor multiple functional outcomes (Fig. 1a). Thus, this ZetaSuite package (outlined in Supplementary Fig. 1b) developed from our splicing screen is generally applicable to other two-dimensional high throughput data.

**Figure 1.**
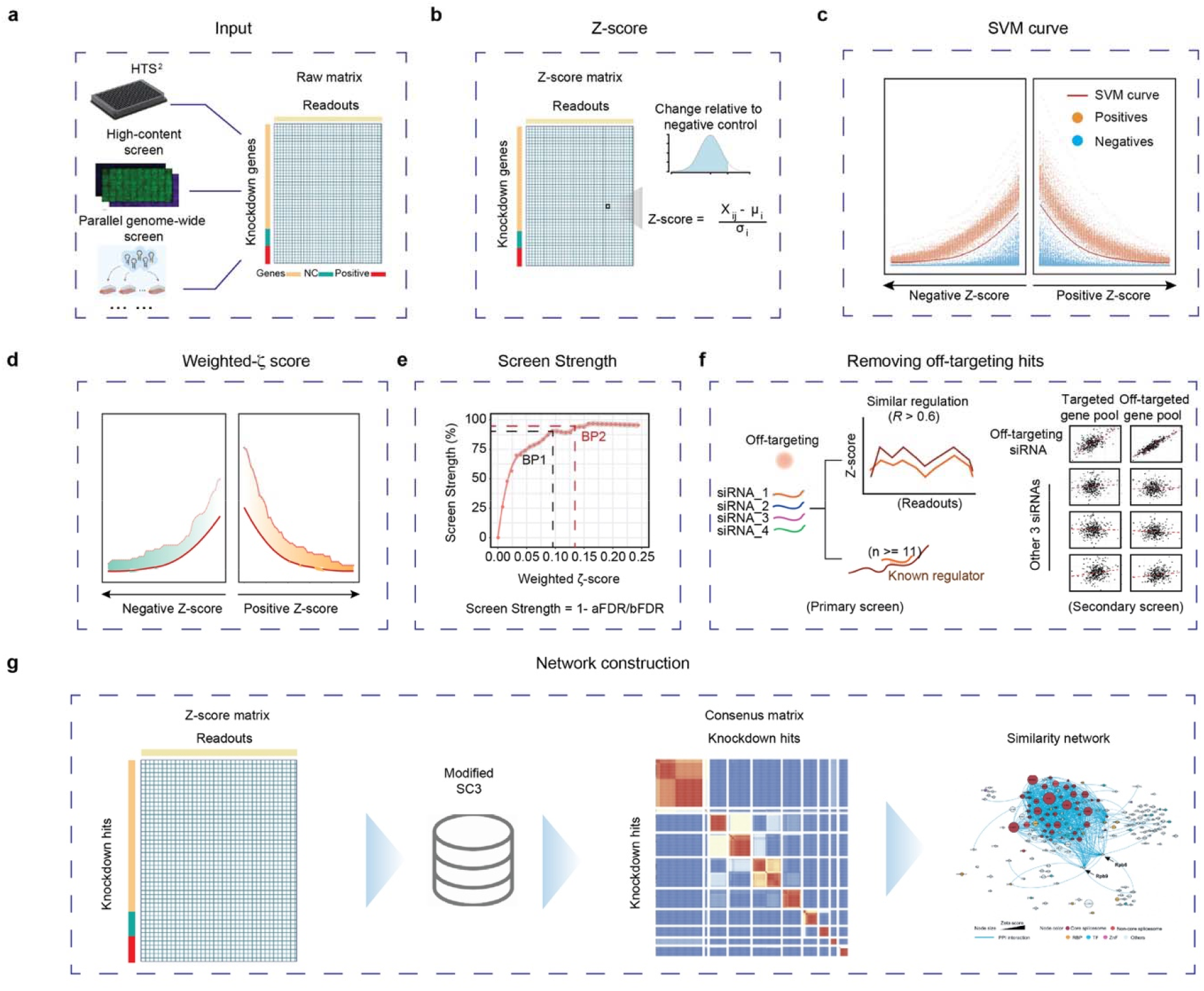
Overview of the ZetaSuite workflow. **a**, Two-dimensional screens include high throughput screen by high through sequencing (HTS^2^), high-content screen, parallel genome-wide screens, etc. ZetaSuite uses the raw matrix as input to calculate ζ score. **b**-**g**, Key steps in the ZetaSuite software from generating initial ζ scores (**b**) to deducing hits by using negative and positive controls to derive a support vector machine (SVM) learning curve (**c**) to calculating weighted ζ scores (**d**) to determining the Screen Strength (**e**) to filtering out off-targets (**f**). The resulting data are used to construct regulatory gene networks based on functional similarities (**g**).

After a series of standard data pre-processing and quality control, ZetaSuite generates a Z-score for each AS event against each targeting RNA (a pool of siRNAs in primary screen or individual siRNAs in secondary screen) in the data matrix (Fig. 1b) and then computes the number of hits at each Z-cutoff from low to high and in both directions to separately quantify induced exon skipping (Fig. 1c, left) or inclusion (Fig. 1d, right) events. This enables the classification of functional data in both directions to identify global splicing activators (if mostly causing exon skipping upon knockdown) or repressors (if mostly inducing exon inclusion events upon knockdown) or both. The same strategy can thus also be used to characterize positive and negative regulators in any specific biological pathway from two-dimensional high throughput data.

When internal positive controls are well separated from negative controls, ZetaSuite calculates a SVM learning curve to maximally separate positives from negatives. Any siRNA that generates a line (a string of data points in the plot) above the SVM line can thus be considered a potential hit and the area between the two line presents the strength of the hit, which can be used to compare and rank hits. We name this statistic as <u>Z</u>-based <u>e</u>stimate of <u>ta</u>rgets or Zeta (ζ) (Fig. 1d). Even without positive controls in certain applications, it is still possible to calculate the area under each line to generate a ζ score for a given hit, which can be used to infer the relative strength of individual hits and rank-order them to deduce important biological information.

As with all screens, a threshold is needed to call hits. To this end, we utilize a large set of non-expressed genes in a given cell type (HeLa cells in our screen) as internal negative controls and determine the number of hits above a given ζ to plot against the number of non-expressed genes mistakenly identified as hits, which may result from either experimental noise or off-target effects. We call this as a Screen Strength (SS) plot, and we select a balance point(s) as threshold where further increase in ζ no longer significantly improves the value of the SS (Fig. 1e). Finally, ZetaSuite takes full advantage of two-dimensional high throughput data to calculate similarities in global responses through pairwise comparisons, which can be leveraged to deduce off-target effects based on the results from the secondary screen (Fig. 1f), and more importantly, to construct gene networks for functional analysis of screen hits (Fig. 1g). Together, ZetaSuite provides a comprehensive package for analyzing two-dimensional high throughput data.

### Increasing readout number leads to diminishing screen specificity with traditional methods

Z or SSMD statistic has been typically used to identify hits from one-dimensional high throughput screens. SSMD has advantages if a screen includes multiple replicates for each targeting RNA ^15^. When the number of screen readouts increases, however, various random outliers become accumulated, which has the potential to severely compromise the screen specificity. For instance, we scored ∼400 AS events against each siRNA and 368 events passed the quality control, and if any of these readouts meets a chosen cutoff, the probability of experimental noise and/or off-target effects would be aggregated in proportion to the number of readouts scored. To demonstrate this, we chose a stringent cutoff of Z>=3 ^24^ to identify hits from our splicing screen data and used siRNAs that target non-expressed genes as true negatives to estimate the screen specificity. Randomly selecting 50 siRNAs against non-expressed genes based on 5 randomly selected AS events, we identified 1 hit out of these 50 true negative siRNAs (Fig. 2a). When all 368 AS events scored in our screen were taken into consideration, the majority of those true negative siRNAs became hits (Fig. 2b). This alarming high false positive rate became further evident when all RNA-seq identified non-expressed genes were included in the analysis (Supplementary Fig. 2a,b). By selecting an increasing number of AS events as readouts to determine the screen specificity, we found that the screen specificity was progressively decreased (Fig. 2c) and we obtained the same result by performing a similar analysis based on SSMD (Fig. 2d). In both analyses, we noted that the specificity was reduced to about half when the number of functional readouts was increased to 50. This illustrated that the most popular statistical approaches for analyzing one-dimensional screen data are no longer suitable for processing two-dimensional high throughput data. Even after using the multiple testing correction methods (such as FDR and Bonferroni correction methods), the number of false positive hits were still very high (see Methods).

**Figure 2.**
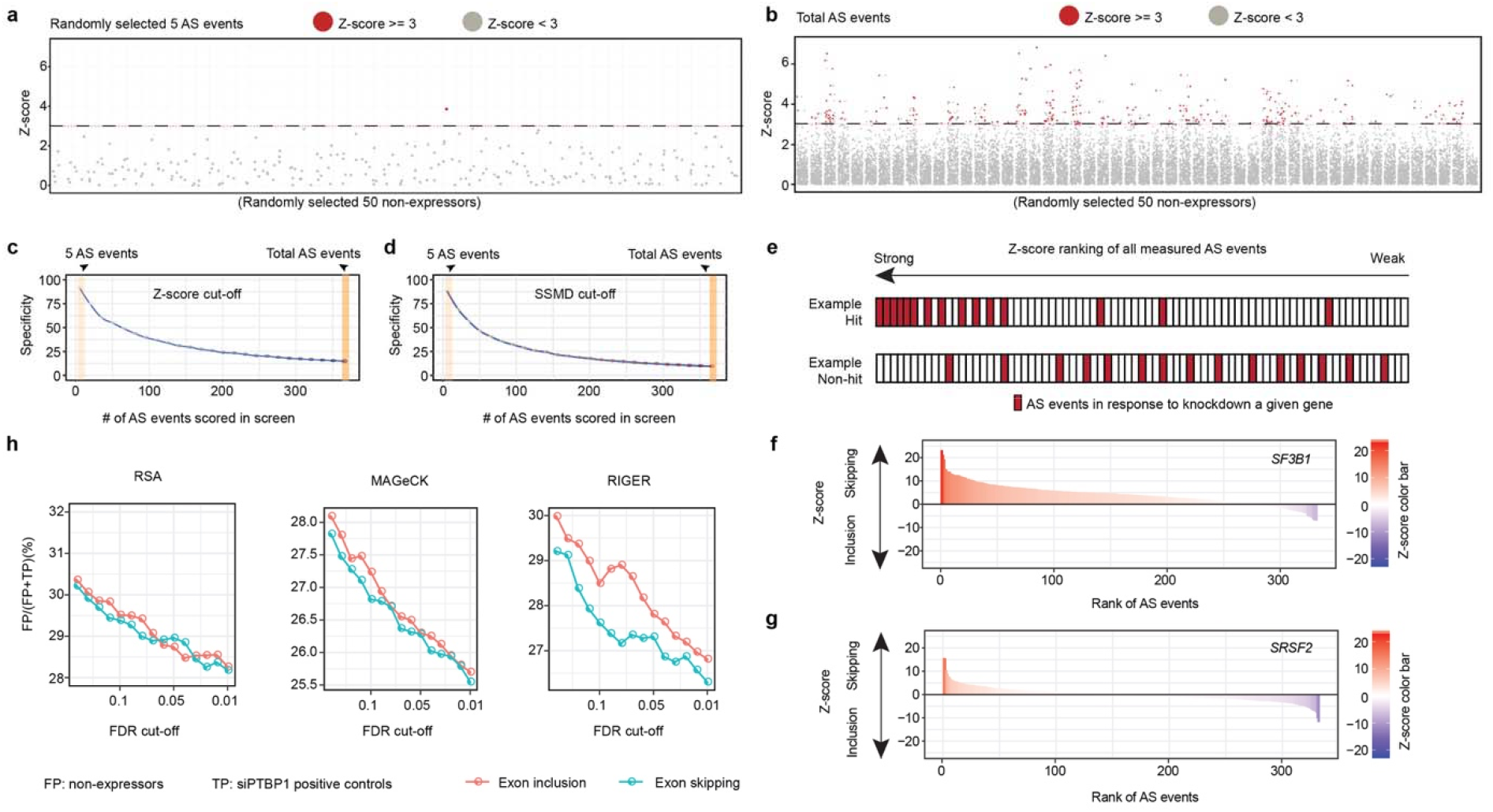
Increasing readout number leads to diminishing screen specificity with common statistical approaches. **a**-**b**, The distribution of Z-scores based on 5 randomly selected alternative splicing (AS) events monitored in our screen (**a**) or all AS events measured (**b**) in response to siRNAs against 50 randomly selected non-expressed genes. The AS event was marked as red if the Z-score is >=3. **c**-**d**, The Specificity based on common cutoffs (**c**, Z>=3) or SSMD (**d**, SSMD>=2) when different numbers of AS events were monitored. The specificity (defined by 1 minus the number of non-expressors scored as hits over the total number of non-expressors) is the mean value of 50 replicates under each condition. **e**, Illustration of the principal theory to determine hits based on RSA, MAGeCK and RIGER. Induced changes in AS are first ranked and the effects of knocking down a given gene on individual AS events are displayed as red bars. A hit would show enriched AS events in one direction (top) while a non-hit would display a relatively random distribution (bottom). **f**-**g**, The distribution of induced AS events (based on Z-scores of induced exon skipping from left to right at top or induced exon inclusion from right to left at bottom) in response to knockdown *SF3B1* (**f**) or *SRSF2* (**g**). **h**, The false discovery rate (FDR=FP/(FP+TP)) at different cutoffs with different methods. The FDRs at x-axis were calculated by different methods (RSA, RIGER and MAGeCK). The FDRs at y-axis were deduced based on the non-expressors and build-in positive controls (siPTBP1). False Positive (FP): non-expressors; True Positive (TP): siPTBP1-treated samples.

We next wondered whether we might adapt the concept from some more sophisticated methods to analyze two-dimensional high throughput data. For example, RSA ^16^, RIGER ^17^, and MAGeCK ^18^ are each designed to determine the impact of a given gene on a functional readout (e.g. cell proliferation) by testing multiple targeting RNAs against each gene and then aggregating the data to reflect the overall contribution of such gene to the functional consequence. A typical data aggregation strategy is analogous to Gene Set Enrichment Analysis (GSEA) ^25^, which is to first rank order all targeting RNAs against all targeted genes tested in a screen based on their functional impact (impact on cell proliferation from left to right) and then score hits if multiple targeting RNAs are enriched at left (Fig. 2e, top raw) whereas a non-hit lacks any enrichment (Fig. 2e, bottom raw).

Here, by replacing individual targeting RNAs with individual AS events, we took a similar strategy to evaluate the overall contribution of a given gene to global splicing control. Using two well-known splicing regulators as benchmarks and separately rank ordering their impact on exon skipping (left to right) or inclusion (right to left), we found that knockdown of the core spliceosome component SF3B1 mainly caused exon skipping (Fig. 2f and Supplementary Fig. 2c), whereas depletion of the SR protein family member SRSF2 induced both exon inclusion and skipping in about equal frequency (Fig. 2g and Supplementary Fig. 2d), consistent with the literature information ^26,27^. Extending this analysis genome-wide, we identified thousands of genes as putative splicing regulators by using different aggregation strategies associated with RSA, RIGER, or MAGeCK (Supplementary Fig. 2e). To evaluate the performance of these methods, we took advantage of 5,006 siRNAs against non-expressed genes as internal negative controls and 299 technical repeats with an siRNA against a well-known splicing regulator PTB as internal positive controls in our screen to estimate the false discovery rate (FDR=false positives divided by false positives plus true positives). We observed an alarmingly high error rate with each of these approaches even at the most stringent FDR cutoff (Fig. 2h). This is likely due to the accumulation of experimental noise plus off-target effects, as illustrated with traditional Z- or SSMD-based approaches (see Fig. 2a,b and Supplementary Fig. 2a,b). These analyses thus present a compelling paradigm for the need to develop a new statistical approach in order to fully explore the power of two-dimensional high throughput data.

### Z-based estimation of global splicing regulators

It becomes quite evident from above analysis that the accumulative experimental noise and off-target effects is a major problem in analyzing two-dimensional high throughput data, such that the screen specificity is progressively diminished as the number of readouts increases. To begin to develop a new statistical strategy to address this problem, we first used non-expressed genes to characterize the distribution of random splicing responses from all AS events quantified in our screen. For each siRNA against a given non-expressed gene, we calculated Z for the entire collection of the AS events scored and then displayed the number of “hits” at each Z-cutoff from low to high for induced exon skipping (toward right) or exon inclusion (toward left). This showed the progressive decline in the number of hits in both directions as Z increases, and after analyzing 10 randomly selected non-expressed genes this way, we noted that all exhibit a similar distribution (Fig. 3a, grey color). In comparison, 10 representative splicing regulators (Supplementary Fig. 3a) all scored a much higher number of hits at any given Z cutoff (Fig. 3a, individually colored).

**Figure 3.**
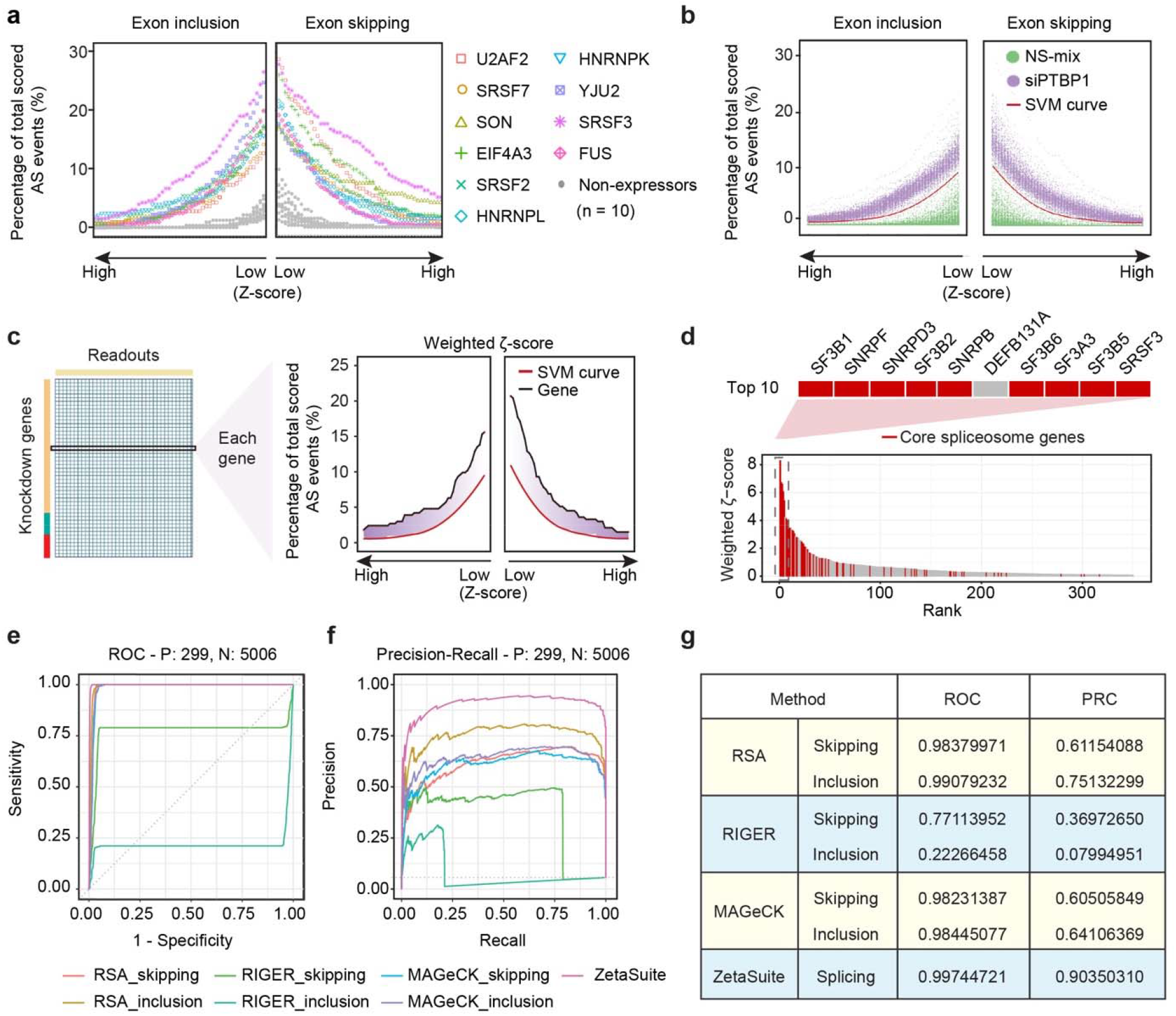
The ζ statistic and comparison with several key existing statistical approaches. **a**, At each Z-score bin over a full Z-range, the level of hits (expressed as the percentage of induced AS events over the total number of AS events monitored) is plotted with 10 representative splicing regulators (individually colored) compared to 10 non-expressors (grey). Left and right separately plot induced exon inclusion and skipping events. **b**, At each Z-score bin over a full Z-range, the level of hits in response to siPTBP1 (purple) or negative controls (NS-mix, green). An optimal SVM curve (red) is derived to maximally distinguish between true positives (siPTBP1) and true-negatives (NS-mix). **c**, Calculation of a weighted ζ-score based on the area between the specific Z-score line of a gene (black) and the SVM curve (red). At each Z-bin, the area is multiplied by the Z-value, thus giving increasingly weights (purple) to hits at higher Z-scores. **d**, The distribution of weighted ζ-score for annotated core spliceosome components among top 350 high-ranking genes. The top 10 high-ranking genes are enlarged (top). Only *DEFB131A* doesn’t belong to core spliceosome, which was later determined to result from off-targeting to *SF3B1* (see Supplementary Fig. 4c). **e**-**f**, The ROC (**e**) and PRC (**f**) curves are deduced using different methods. Weighted ζ-score in two directions calculated by ZetaSuite are combined in this analysis to reflect the overall functional consequence. This is not applicable to other methods, and we thus display the data separately. **g**, The summary of the areas under all deduced ROC and PRC curves using different methods.

Interestingly, such distinct profiles between non-expressors and known splicing regulators were similarly observed with a large number of built-in negative controls with a pool of non-specific siRNAs (NS-mix) and positive controls with a specific siRNA pool against *PTBP1* (siPTBP1). This enabled us to develop a SVM curve to maximally separate positives from negatives (Fig. 3b). We define the area between a putative hit above the SVM line as a <u>Z</u>-based <u>e</u>stimate of <u>ta</u>rgets or Zeta (ζ). To favor the differences at higher Z cutoffs, we multiply the number of Z with the area at the corresponding range and then aggregate the values from the full range of Z to obtain a weighted-ζ score to define the overall impact of a putative splicing regulator (Fig. 3c), which can be used to rank individual splicing regulators. To characterize a given splicing regulator in splicing activation and repression, we separately calculated its ζ scores for aggregated exon inclusion or skipping events. After processing our splicing screen data with this analysis pipeline (called ZetaSuite, see Supplementary Fig. 1b) and rank ordered the hits according to their overall impact on AS (high to low from left to right), it became evident that most high-ranking hits correspond to annotated core spliceosome components (Fig. 3d). This demonstrated for the first time that components of the core splicing machinery also function as the most prevalent class of AS regulators in mammalian cells. Moreover, these genes are in general highly expressed in mammalian cells and their inactivation predominantly induce exon skipping (Supplementary Fig. 3b,c).

To compare the performance of Zeta with other ranking approaches, such as that used in RSA, RIGER, or MAGeCK, we again took advantage of a large number of built-in positive and internal negative controls in our screen, which allowed us to precisely determine the numbers of true and false positives and negatives to construct Receiver Operating Characteristic (ROC) (Fig. 3e) and Precision-Recall curves (PRC) (Fig. 3f). These comparisons demonstrate that the newly developed ζ statistic significantly outperform all other ranking methods in analyzing two-dimensional high throughput splicing screen data (Fig. 3g).

### Hit selection based on reflection points and Screen Strength

Any screen requires a cutoff to maximize true positives while minimize true negatives. In most one-dimensional high throughput screens, hits are first ranked based on Z or SSMD and the threshold is then determined by estimating the false positive level (FPL) and the false negative level (FNL) ^28^. As Z or SSMD increases, FPL will gradually decrease while FNL will progressively increase. This approach can be similarly applied to ζ-based scoring, as illustrated with our splicing screen data using the siPTBP1 in technical repeats as true positives and siRNAs against non-expressed genes as true negatives (Supplementary Fig. 4a). Using the balanced error level approach as recommended earlier ^28^, we obtained 0.7% for both FPL and FNL with calculated FDR as 12.3%.

However, many siRNA screens may not be able to build in a large number of true positive controls, and additionally, the balanced error level is likely influenced by how well positive controls can be differentiated from negative controls. We therefore sought to determine a cutoff by only using non-expressed genes as negative controls, which would be generally applicable to most genome-wide screens by RNAi. For this purpose, we introduce the concept of apparent FDR (aFDR), which is defined as the number of non-expressors identified as false positive hits among all hits scored at a given cutoff. Before screen, we have a baseline FDR (bFDR), which corresponds to the number of non-expressors among the total number of genes targeted in the screen. By definition, bFDR represents the chance from a random draw. We next define the Screen Strength: SS=1-aFDR/bFDR, which can be used to evaluate the effectiveness a screen has achieved relative to random draw. We can also use this parameter to compare between different screen results. Using this approach, we plotted the SS based on our splicing screen data against increasing ζ (Fig. 4a). This allows us to calculate the balance point (BP) for hits selection where the SS will almost no change as the stringency increases. With our splicing screen data, we actually identified two such BPs, thereby enabling us to define candidate hits after BP1 and high confidence hits after BP2, the latter of which maximally eliminate true false positives derived from non-expressors (Fig. 4b).

**Figure 4.**
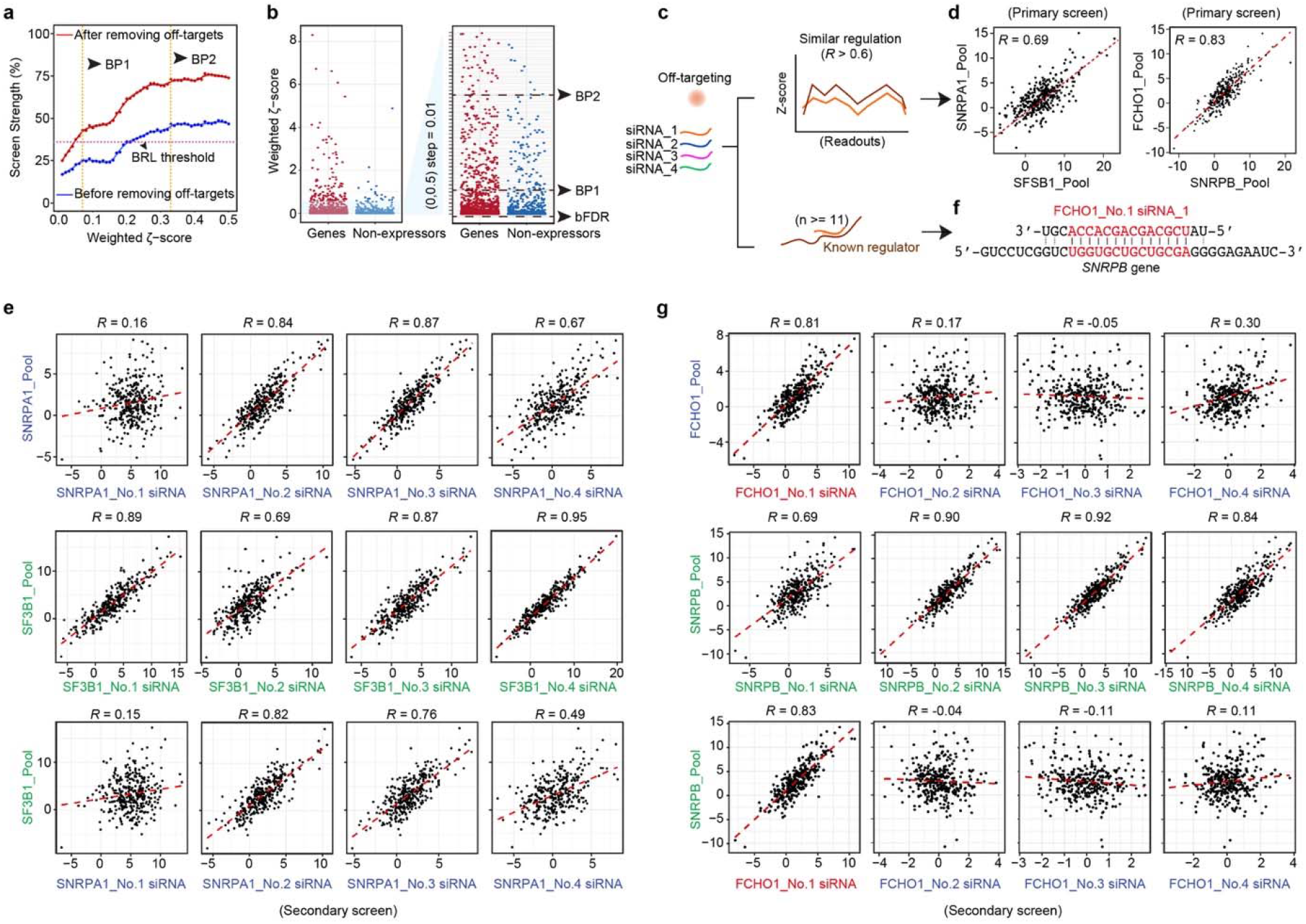
Hit selection based on Screen Strength and strategy to filter out off-target effects. **a**, The comparison of the Screen Strength before (blue) and after (red) filtering out off-targets. BP: balance point. Note that the Screen Strength based on the threshold defined by the commonly used balanced error level (BRL) approach is also indicated (see Supplementary Fig. 4a). Those between BP1 and BP2 are candidate hits and those after BP2 are high confidence hits. **b**, Weighed ζ-scores of expressed and non-expressed genes. A specific region is enlarged on the right for comparative purpose. bFDR: baseline FDR. BP1 and BP2 are according to those defined in **a**. **c**, Strategy to filter out off-target effects based on similarity in response and sequence complementarity. **d**, Comparison of AS events responsive to knockdown of *SNRPA1* and *SF3B1* or *SNRPB* and *FCHO1* in primary screen. Pearson correlation score is indicated in each case. **e**, Comparison of AS events responsive to knockdown of the siRNA pool vs individual siRNAs against *SNRPA1* or *SF3B1* in the secondary screen. The third row shows the comparison between the siRNA pool against *SF3B1* and individual siRNAs against *SNRPA1*. **f**, The sequence of a single siRNA targeting *FCHO1* is aligned with its potential off-target on the *SNRPB* transcript. **g**, Comparison of AS events responsive to knockdown of the siRNA pool vs individual siRNAs against *FCHO1* or *SNRPB* in the secondary screen. The third row shows the comparison between the siRNA pool against *SNRPB* and individual siRNAs against *FCHO1*. Red highlights the predicted off-targeting siRNA.

### A strategy to remove off-target effects from two-dimensional high throughput screen data

Off-target effects have been a major problem in genome-wide screens. Recent strategies to filter out off-targeting RNAs are to increase the number of targeting RNAs against each gene and eliminate those that show divergent effects from the consensus generated by multiple targeting RNAs ^19^. These approaches are based on the assumption that an activity defined by the majority of targeting RNAs reflects on-target effects, which may not always be the case. In addition, these approaches require a large number (usually 15 to 20) of targeting RNAs per gene, thus unapplicable to traditional of siRNA or shRNA libraries that typically contain 4 to 6 targeting RNA in each pool. In fact, the increased sequence complexity may also cause additional off-target effects. We thus sought to utilize the data from primary and secondary screens with traditional arrayed siRNAs to filter out off-targets, again by taking advantage of multiple functional readouts from each treatment.

As illustrated in Fig. 4c, we first identify siRNA pools that show similar responses in pairwise comparison, which we define by requiring R>=0.6 ^29^. Because two genes may have related function in a common biological pathway, more than one siRNA in their pools often show similar responses to both of their pools in the secondary screen, as illustrated with *SNRPA1* and *SF3B1*, both being subunits of the U2 ribonucleoprotein particle (snRNP) (Fig. 4d and 4e). This is further illustrated with multiple core spliceosome components (Supplementary Fig. 4b). On the other hand, when a similar response results from some off-targeting effects, we found in most cases where one specific siRNA in a given siRNA pool shows sequence complementarity of consecutive 11nt or longer to the transcript targeted by the other siRNA pool (see Fig. 4f), as shown earlier when examining cross reacting siRNAs ^30^. Furthermore, it is this same siRNA that is also responsible the similar response in the secondary screen, as exemplified with *FCHO1* and *SNRPB* (Fig. 4g). Indeed, in this case, *SNRPB* is a known core spliceosome component, whereas *FCHO1* is a gene functioning in early step of clathrin-mediated endocytosis ^31^, but without any documented role in regulated splicing, suggesting that the high ζ value generated by siFCHO1 resulted from its off-target effect on *SNRPB*. We thus propose a general strategy to eliminate potential off-target effects if a single siRNA in a given pool is responsible for (i) generating a similar functional response and (ii) shows a significant sequence complementarity to the transcript targeted by another siRNA pool. Using this strategy, we identified multiple siRNA pools that likely caused off-targets due to specific cross reactions with well-established splicing regulators (Supplementary Fig. 4c).

We extended this analysis to all non-expressors in our screen and showed that filtering out those with identifiable off-targeting activities significantly improved the Screen Strength (Fig. 4a, from blue to red line). Moreover, ζ scores may differ when different positive controls are used to generate the SMV. To evaluate this impact, we focused on high confidence hits after BP2 based on using repetitive siPTBP1 treatments as positive controls and found that >90% of hits were identifiable with a different set of internal positive controls (see Supplementary Fig. 3a) to deduce a slightly different SVM line (Supplementary Fig. 4d,f), suggesting that slightly distinct positive controls only affect low-ranking candidates. Because of the ability to rank the hits, we were also able to detect >90% of the hits based on siPTBP1-derived SVM based on the balance point alone without using any SVM (Supplementary Fig. 4e,f), although the ability to generate a SVM curve helps minimize inclusion of low confidence hits, which moderately affected multiple readouts. Finally, we evaluated the performance of the ζ statistic on different numbers of functional readouts. Using true positives (siPTBP1) and high confidence hits based on using all AS readouts as the reference sets, we tested whether ζ was able to detect these ’reference’ genes using fewer readouts and found that ζ was able to identify over 80% of these ’reference’ genes when the readout size reaches 200 or greater (Supplementary Fig. 4g). This information offers a general guide to designing future two-dimensional genome-wide screens.

### Application of ZetaSuite to understand core fitness genes in cancer cells

Having established the general framework for the ζ statistic with our own splicing screen data, we next sought to test its general applicability to other large-scale data resources. DRIVE ^10^ and DepMap ^11^ are representative of such data, both designed to determine cancer dependencies by transducing a large panel of cancer cell lines with pooled shRNAs and identify depleted shRNAs by deep sequencing to quantitatively report the essential function of their targeted genes in individual cancer cell lines. DRIVE tested more cell lines than DepMap (overlap=113, Supplementary Fig. 5a), whereas DepMap had more genes covered in its shRNA pools than DRIVE (overlap=7,081, Supplementary Fig. 5b). Thus, similar to our splicing screen data set, the first dimension consists of individual RNAi treatments and the second corresponds to multiple functional readouts (different AS events vs different cell lines). Additionally, similar to our experimental design, DepMap selected a set of known essential genes (n=210) ^20^ as positive controls and used non-expressed genes (n=855) as negative controls. We found that these controls are well separated based on t-distributed stochastic neighbor embedding (tSNE) ^32^ with both data sets (Supplementary Fig. 5c).

For data analysis, DRIVE utilized RSA to rank order hits and ATARiS to eliminate shRNAs that may cause off-target effects. A gene was considered essential if RSA>= -3 in >50% of the cell lines tested. In contrast, DepMap removed off-target effects with DEMETER and selected top hits showing 6 standard deviation (SD or σ) or greater in any cell line tested for further pathway analysis. As we demonstrated in treating our two-dimensional splicing screen data, an arbitrary cutoff would always present a trade-off between sensitivity and specificity, and even with the most extreme cutoff like 6σ, experimental noise would still become accumulated with the increasing number of readouts scored in a screen. We thus introduced a general parameter in ZetaSuite, the Screen Strength (SS), to compare between different screen results.

Here, we processed the data from DepMap and DRIVE with the ZetaSuite pipeline (see Supplementary Fig. 1b). Although DRIVE and DepMap mainly determined cancer dependencies by scoring depleted shRNAs, we wondered whether the data sets also contain useful information on enriched shRNAs, which would be indicative of some opposite functions to cancer dependency, referring here to as cancer checkpoint. To simplify the comparison between the two data sets, we chose to start with the processed data with potential off-target effects already removed to quantify depleted and enriched shRNAs. We then plotted the DepMap and DRIVE data in both directions in the full range of cut-offs. As expected, positive controls and non-expressors are well separated in both data sets in the direction of cancer dependency (Fig. 5a). We next calculated the weighted ζ-score for each tested gene in both data sets and then displayed the data in the Screen Strength plot (Fig. 5b), from which we determined two balance points (BP1 and BP2) for cancer dependency in both data sets. To identify genes involved in cancer checkpoint, we unable to derive any balance point with the DepMap dataset, likely due to scattered data from a relatively smaller number of cell lines surveyed (Fig. 5b), and for DRIVE, we only used the most stringent cutoff at BP2 to select hits (Fig. 5c).

**Figure 5.**
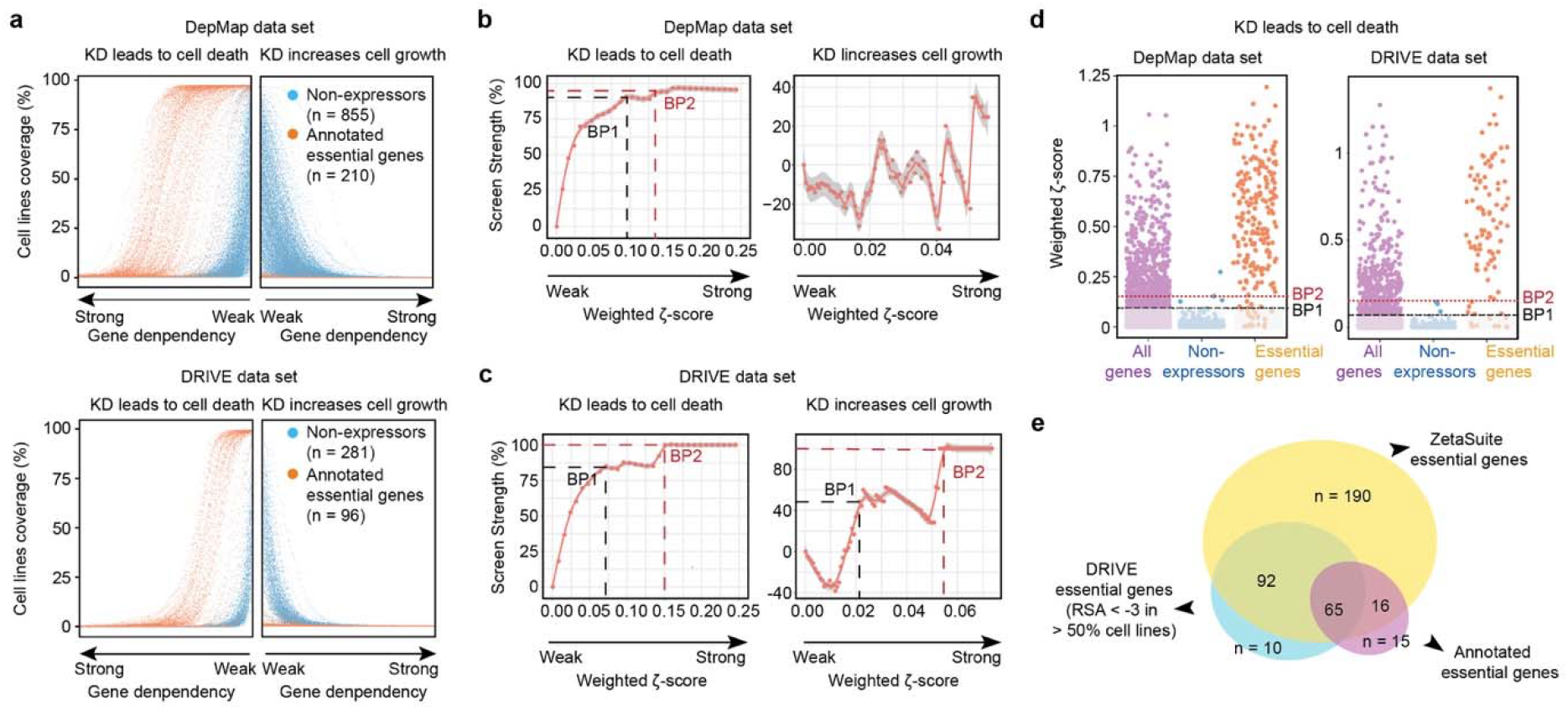
Application of ZetaSuite to mine core fitness genes in cancer cells. **a**, At each gene dependency bin over a full range of gene dependency scores, the percentage of cell lines responsive to knockdown of individual annotated essential genes (orange dots) or non-expressed genes (blue dots) based on the DepMap (top) and DRIVE (bottom) datasets. **b**-**c**, Screen Strength plot at different cutoffs for cancer dependency (left) or cancer checkpoint (right) deduced from the DepMap (**b**) or DRIVE (**c**) dataset. Because of scattered data, balance point could not be determined in the DepMap dataset. The two balance points (BP1 and BP2) in the DRIVE dataset are marked (**c**). **d**, Hits for cancer dependency above the threshold defined by BP1 or BP2 based on the data from DepMap (left) or DRIVE (right). **e**, Comparison of cancer dependencies deduced in the DRIVE project with those newly determined with ZetaSuite and previously annotated essential genes.

Based on the selected BP1 and BP2, we found that the majority of positive controls were included in both data sets, suggesting that ZetaSuite-suggested cutoffs are able to encompass the majority of cancer dependencies, even at BP2 (Fig. 5d). Since DepMap only focused on specific cancer dependencies by requiring 6σ, we compared ZetaSuite-identified hits with DRIVE-defined essential genes and previously annotated essential genes ^33^, and based on BP2, we found that ZetaSuite identified more hits than previous analyses (Fig. 5e). Moreover, none of those 10 DRIVE hits (Fig. 5e, blue) missed by ZetaSuite are part of the annotated essential genes. Despite the significantly enlarged hit size, enriched Gene Ontology (GO) terms, KEGG pathways and Complexes annotated in the CORUM database ^34^ associated with newly identified hits are similar to those deduced earlier based on much more stringent cutoffs, with top ranked terms linked to key housekeeping activities, such as DNA replication, Splicing, Cell cycle, RNA transport, Ribosome biogenesis, etc. (Supplementary Fig. 5d,e,f). Additionally, we noted that these newly identified hits were largely anti-correlated with AGO2 expression and copy number variation (CNV) (Supplementary Fig. 5g), as reported earlier with the DRIVE dataset ^10^. Conversely, 8 out of 10 hits identified by DRIVE but missed with ZetaSuite lack the anti-correlation with both AGO2 expression (Supplementary Fig. 5g, left) and AGO2 CNV (Supplementary Fig. 5g, right). Together, these data demonstrated the effectiveness and objectiveness of ZetaSuite in identifying cancer dependencies from previous large-scale screen data.

### Biological insights into cancer dependency

The expanded list of cancer dependencies enabled us to gain further insights into critical cancer development pathways compared to those already recognized from previous analysis with the limited set of genes. For example, based on similarities among different DRIVE cancer cells, we were able to deduce 7 clusters by t-SNE plotting and draw the global network based on regulation similarity for total hits that passed the BP1 threshold (Fig. 6a). One of these gene networks is enriched in components of the transcription mediator complex and Pol II, all connected to the well-known oncogene *MYC* (Fig. 6b), consistent with the known function of MYC in transcriptional control ^35^. Interestingly, MYC inhibition showed the most dramatic impact on rhabdoid cancer cells (Supplementary Fig. 6a), which is in agreement with a recent observation that MYC inhibition effectively restricted rhabdoid tumor growth in vivo ^36^. In this MYC dependency plot, we also noted significant *MYC* dependency in multiple myeloma (MM) cancer cells, which is in line with frequent 8q24 translocation that leads to MYC overexpression in MM cancers ^37^.

**Figure 6.**
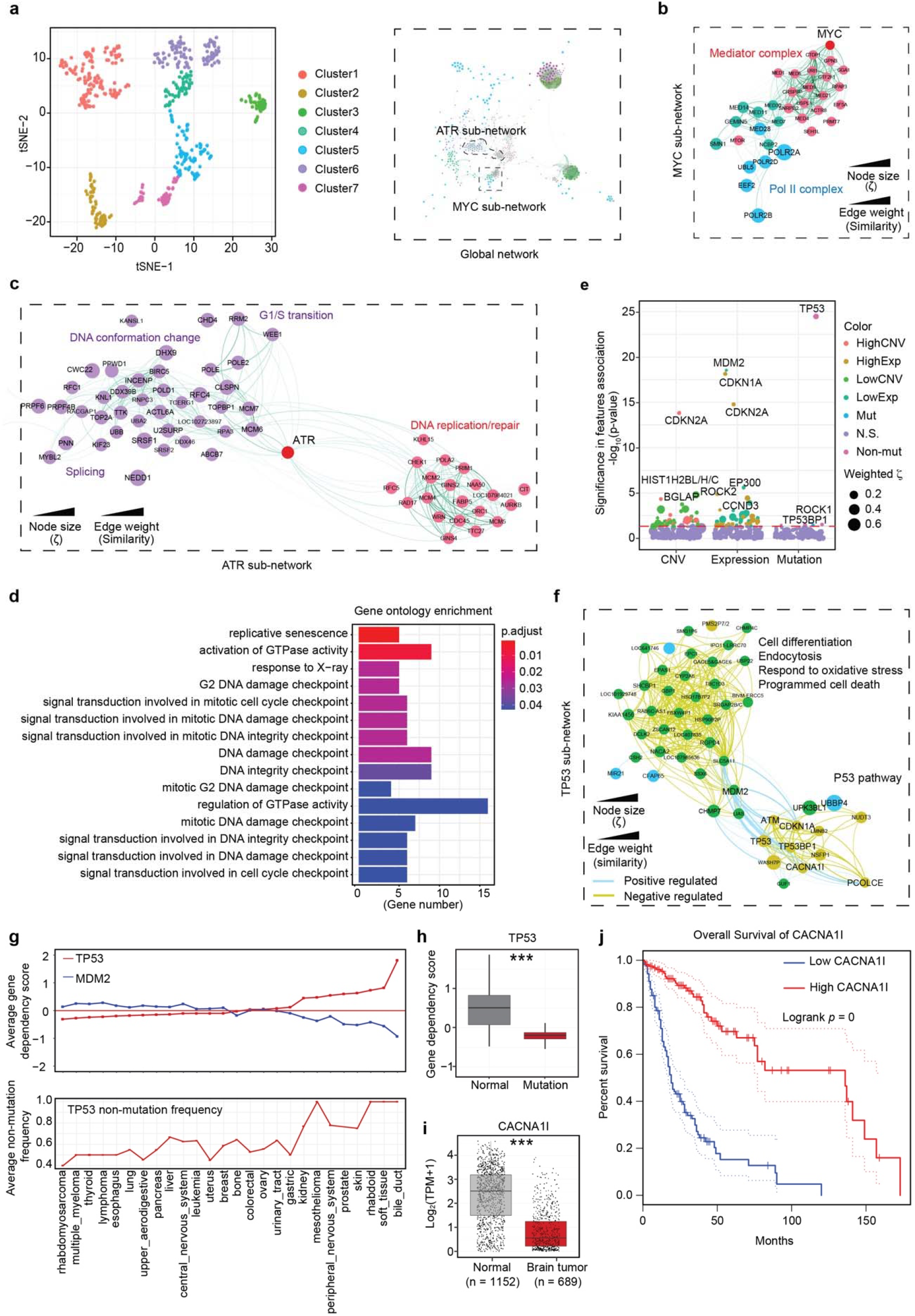
Biological insights from identified cancer dependencies. **a**, Cluster (left) and global network (right) for cancer dependencies determined by ZetaSuite from the DRIVE dataset. **b**-**c**, *MYC*-associated sub-network, highlighting its connectivity to Mediators and Pol II components (**b**), and *ATR* connectivity to sub-networks associated with genes involved in DNA conformation or DNA replication/repair (**c**). Colors correspond to different clusters defined in **a**. **d**, Functionally enriched GO term biology pathways for cancer checkpoint hits based on the DRIVE dataset. Shown are top 15 GO terms with smallest adjust *p*-values. **e**, The association of ZetaSuite-identified cancer dependencies with gene expression, copy number and mutation features. For each gene, cancer cell lines were firstly ranked based on the levels of CNV or gene expression, and the cancer dependency scores were then compared between cell lines in top 25% versus bottom 25%. The *p*-value (y-axis) for each gene in this comparison was determined by Wilcox-test. In addition, for association analysis with mutations, cancer cell lines were divided in two groups with or without mutation for each gene, The cancer dependency scores were then compared between these two groups and the *p*-value (y-axis) in this comparison was determined by Wilcox-test. Some representative genes are highlighted in each feature group. Genes above the red dashed line have *p*-values<0.05. **f**, *TP53*-associated sub-network. **g**, Averaged dependency scores for *TP53* and *MDM2* (top) and *TP53* non-mutation frequency (bottom) in different cancer tissues. Tissues are ranked based on averaged *TP53* dependency scores. **h**, The *TP53* gene dependencies in normal or mutated *TP53* cell lines. *** *p*<0.001 based on Wilcox-test. **i**, *CACNA1I* gene expression in normal brain tissues (based on the GTEx database) and brain tumors (based on the TCGA database). *** *p*<0.001 based on Wilcox-test. **j**, Kaplan-Meier survival curves of brain tumor patients associated with high or low *CACNA1I* expression. The dashed lines indicate the 95% confidence intervals.

We also detected two separate clusters connected by *ATR*, a key regulator of genotoxic stress. One cluster includes various genes involved in G1/S transition and modulation of DNA topology and the other encompasses genes critical for DNA replication/repair (Fig. 6c). This is consistent with the existing literature on the function of ATR in connecting genotoxic stress to cell cycle control ^38^. Interestingly, we noted several splicing regulators (i.e., *SRSF1* and *SRSF2*) in these clusters, both being previously implicated in inducing aberrant R loops that led to ATR activation ^39^. This has been suggested as a key mechanism underlying Myelodysplastic Syndromes (MDS), a pre-leukemia that can rapidly progress to acute myeloid leukemia (AML), thus explaining greater ATR dependency in leukemia than most other cancer types (Supplementary Fig. 6b). These data further demonstrated the utility of the ZetaSuite in analyzing the DRIVE and DepMap datasets to mine important cancer pathways.

### Genes involved in global cancer checkpoint

One of the most significant advances in further mining the DRIVE dataset with ZetaSuite is the discovery of genes whose depletion appears to promote tumor growth (i.e., those deduced from enriched shRNAs). Strikingly, GO term analysis of these genes revealed that the vast majority of them are involved in DNA checkpoint control (Fig. 6d). Previously, genes involved in cancer dependencies were cross analyzed with copy number variation (CNV), gene expression, or mutation frequencies, revealing their association with low CNV and low expression, which has been referred to as CYCLOPS genes ^40^. We also confirmed this finding with ZetaSuite-identified cancer dependencies (Supplementary Fig. 6c). We performed a similar analysis on cancer checkpoint genes and identified 9 major clusters (Fig. 6e). Contrary to core fitness genes, much fewer cancer checkpoint genes were associated with CNV, altered expression, or mutation in DRIVE cell lines.

Several typical tumor suppressors were identified as strong cancer checkpoints in this feature association analysis, including *TP53* (encoding for p53) ^41^ and its transcription target *CDKN2A* (encoding for the cell cycle inhibitor p16) ^42^ and *CDKN1A* (encoding for the cell cycle inhibitor p21) ^43^. Interestingly, *MDM2*, an E3 ligase for p53, was also identified as a cancer checkpoint gene (Fig. 6e). The similarity network clearly reflected the antagonizing function between *TP53* and *MDM2* (Fig. 6f). In fact, while wildtype *TP53* always gave rise to a positive dependency score, reflecting its tumor suppressor function, mutant *TP53* produced a negative cancer dependency score, indicating its oncogenic role in those tumor cells (Fig. 6g,h), which is in full agreement with the known functions of wildtype and mutant p53 in tumorigenesis ^44^. Most interestingly, as exemplified with *MDM2*, we observed that multiple cancer checkpoint genes are also linked to either low CNV or low expression (see Fig. 6e), suggesting that the CYCLOPS phenomenon also applies to some key cancer checkpoints. *MDM2* was also connected to a cluster of genes functioning in cell differentiation, endocytosis, cell death and response to oxidative stress, consistent with the role of MDM2 in regulating the transition from proliferation to differentiation ^45^ and in the cellular response to oxidative stress ^46^.

In the elucidated p53 subnetwork, *TP53BP1* and *ATM* activate *TP53*, which in turn activates *CDNK1A* (Fig. 6f). Besides these known functional connections, we also identified various genes without prior connection to the p53 pathway, such as *PCOLCE* and *CACNA1I*. As an extracellular matrix protein and a major regulator of fibrillar collagen biosynthesis, disruption of *PCOLCE* has been reported to induce cell growth in cultured fibroblasts, suggesting a role in cell proliferation control ^47^. *CACNA1I*, a gene involving controlling voltage-gated calcium channels, was significantly down regulated in brain tumors compared to surrounding normal tissues (Fig. 6i) and patients with low *CACNA1I* expression were associated with poor prognosis based on the TCGA database (Fig. 6j). The newly discovered connection of this and other critical genes with the p53 pathway will fuel future studies on tumorigenesis.

Last, but not least, further analysis of the newly identified cancer checkpoints revealed several major regulatory gene networks based on their similarities among different DRIVE cell lines (Supplementary Fig. 6e). Besides those critical for cell aging, such as *TP53*, *CDKN2A*, *BGLAP*, *CDKN1A*, as described above, we also noted gene networks for phosphorylation regulation (e.g., *MAP3K9*, *TAOK1*, *ROCK1/2*), GTPase activities (e.g., *EPHA5*, *TBC1D3D*, *RND3*), and DNA packaging (e.g., *HIST1H2BN*, *HIST1H2BL/H/C*). These findings not only support the documented roles of specific MAPK and Rho GTPase pathways in coordinating regulated tumor growth ^48,49^, but also raise a new paradigm regarding how DNA packaging proteins may promote tumor growth. Collectively, this functional connectivity map provides critical insights into the involvement of an elaborated gene network in checkpoint control, which may be critical for long-term cell survival, even among cancer cells.

## Discussion

The increasing power and decreasing cost with deep sequencing technologies have enabled multi-dimensional analyses of gene expression. By coupling high throughput screening with high throughput sequencing (HTS^2^), it is possible to utilize a specific set of genes as a surrogate for specific cellular activities in chemical and genomic screens ^8,9^. By monitoring hundred or even thousand functional readouts, such “ultrahigh-content” screens offer numerous advantages over traditional one-dimensional screens, among which include the ability to deduce gene networks directly from the primary screen results and the feasibility to perform a drug screen without relying on a pre-defined druggable target ^8,9^. More recently, we have extended the HTS^2^ approach to a genome-wide screen by scoring hundreds of alternative splicing events to identify global splicing regulators ^23^, illustrating a two-dimensional screen strategy that can be adapted to study many different paradigms in regulated gene expression.

This added dimension also requires a coordinated effort in developing a suitable statistical model for data analysis. We therefore developed a new ζ statistic, and using our in-house HTS^2^ data on global splicing regulators, we demonstrated that ζ outperforms all existing strategies based on hit ranking and aggregation, such as RSA ^16^, RIGER ^17^ and MAGeCK ^18^, all of which are based on the null hypothesis that most of screened genes are non-hits. These existing methods are thus not suitable for analyzing data from secondary screen or using pre-selected candidates. In contrast, the ζ statistic can be broadly used to process two-dimensional data, which only requires a large number of negative controls. Interestingly, as demonstrated in the current study, non-expressed genes can be considered a large set of internal negative controls. In ZetaSuite, we also introduce the Screen Strength to measure the success of a given screen and compare between screens.

Off-target effects represent a major problem in genome-wide screen with siRNAs, shRNAs, or sgRNAs. To reduce the impact of off-target effects, one strategy is to increase the number to targeting RNAs (up to 50 per gene) to target each gene ^50^. Multiple algorithms have been developed to remove potential off-target effects. For example, ATARiS was developed based on the assumption that multiple on-targeting RNAs would give rise to similar results while off-targeting RNAs would each cause a distinct non-specific effect ^19^. This assumption has the potential to retain off-targeting hits if multiple targeting RNAs cause similar non-specific effects, for instance, due to commonly induced cellular stress. In comparison, DEMETER ^11^ or its recently refined version DEMETER 2 ^20^ filter out off-targeting effects based on the assumption that off-targets likely result from the sequences in the “seed” region to cause microRNA-like effects on other genes. This assumption may not be reliable because of the very relax “seed rule” and various miRNA-like effects induced by sequences outside the seed region ^21^. In contrast to these existing approaches, ZetaSuite eliminates off-targets based on two criteria, one on the functional similarity and the other on the sequence complementarity between a targeting RNA and an off-targeted transcript. Interestingly, by leveraging the results from the secondary screen, we found that a single siRNA in a pool is often responsible for the off-targeting effect of that pool and the same siRNA also shows the complementary sequence to the predicted off-target. Our strategy thus enables the utilization of traditionally designed arrayed libraries for two-dimensional genome-wide screens.

After demonstrating the power of the new ζ statistic to rank order global splicing regulators and revealing a predominant role of the core splicing machinery in the regulation of alternative splicing, we further took the ZetaSuite to re-analyze the large-scale data from public DRIVE and DepMap cancer dependency projects, which were designed to tackle cancer dependencies. Interestingly, prior efforts in analyzing these datasets have been primarily focused on cancer dependencies, revealing various gene networks critical for cancer cell survival. DRIVE defines cancer dependency by requiring RSA>= –3 on >50% of cell lines surveyed while DepMap paid attention on specifically regulated hits with 6σ or greater. These definitions appear to be relatively arbitrary and unnecessarily too stringent without fully exploring the power of such large-scale datasets. By revisiting these data with the newly developed ZetaSuite, we have now elevated cancer dependencies by several folds, leading to the elucidation of several new critical gene networks contributed by some well-established oncogenes and tumor suppressors, such as *MYC*, *ATR*, and *TP53*. These discoveries potentiate further dissection of these fundamental oncogenic pathways.

Perhaps the most intriguing observation from analyzing the DRIVE dataset is the identification of genes whose depletion appears to accelerate cancer cell proliferation, at least transiently during the treatment period. Strikingly, the vast majority of these hits function in various DNA checkpoint pathways, which we refer to as cancer checkpoint, opposite to cancer dependency. Interestingly, we note that some of those cancer checkpoints are also linked to CNV and low expression, and although less prevalent compared to cancer dependencies, this observation suggests that genes involved cancer checkpoints are also related to the phenomenon of CYCLOPS ^40^. Such depletion-induced cell proliferation may allow cancer cells to temporally escape DNA checkpoint control, indicating that various cancer cells still retain such programs to protect their unstable genomes from becoming further deteriorated. In this regard, the exposure of these new cancer vulnerabilities may aid in the development of new cancer therapies, as exemplified by using ATR inhibitor to treat MDS ^51^.

## Methods

ZetaSuite is designed to address challenges in analyzing two-dimensional high throughput data. Supplementary Fig. 1b provides an overview of the flow chart, as individually detailed below.

### ZetaSuite Part 1

#### Data preprocessing

Before running the main ZetaSuite procedure, raw data are first filtered to remove low-quality samples (columns) and readouts (rows) in the data matrix to minimize false positives. The default threshold is set to remove a row or a column if the number of drop-outs (missing values; in our study, a value is missing if one of the mRNA isoforms is undetectable) is larger than the value of Q_3_+3*(Q_3_-Q_1_) where Q_1_ and Q_3_ are lower and upper quartile, respectively. The remaining data are processed with the KNN-based method to estimate the missing values with the parameter k=10.

### ZetaSuite Part 2

#### QC evaluation

Quality Control (QC) is a critical step in evaluating the experiment design. For all two-dimension high throughput data, t-SNE plot ^32^ is first used to evaluate whether features are sufficient to separate positive and negative controls. The SSMD score ^14^ is further generated for each readout to evaluate the percentage of high-quality readouts. In our case, the data will be further processed if >5% of reads are of the SSMD score >2.

#### Conversion of input matrix to Z-score matrix

After data pre-processing, the initial input matrix is arranged in N x M dimension, where each row contains individual functional readouts against a siRNA pool and each column corresponds to individually siRNA pools tested on a given functional readout. Readouts in each column may be thus considered as the data from one-dimensional screen (many-to-one), and thus, the typical Z statistic can be used to evaluate the relative function of individual genes in such column. The conversion is repeated on all columns, thereby converting the raw activity matrix into a Z-score matrix. Suppose N_ij_ are the values in the original matrix i (1≤ i ≤ N siRNA pool) row and j (1≤ j ≤ M readout) column, then

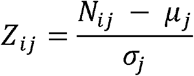

Where *μ_j_* and *σ_j_* are the mean and standard deviation of negative control samples in column j.

#### Generation of Zeta plot

The x-axis in the Zeta plot shows a series of Z-score cutoffs in two directions (in our case, induced exon skipping in the positive direction and inclusion in the negative direction) and the y-axis is the percentage of readouts survived at a given Z-score cutoff over the total scored readouts.

To generate this plot, the range of Z-scores is first determined by ranking the absolute value of total Z_ij_ (Z-score value in row i and column j) from the smallest to the largest (|*Z*_1_|, |*Z*_2_|, …|*Z_i_*_−1_|, |*Z_i_*|, |*Z*_*i*+1_|, …,|*Z_N × M_*|, where |*Z*_*i*−1_|≤|*Z_i_*|≤|*Z*_*i*+1_| nd i here is the rank number). To exclude insignificant changes that may result from experimental noise (|Z|<2, which equals to *p*-value>0.05), Z-cutoffs are selected in the range of 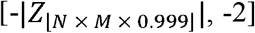 in the negative direction and 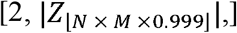 in the positive direction. The Z range in both directions is next divided into 100 bins (B = (*b*_1_, *b*_2_, …, *b_i_*,…, *b*_100_), where 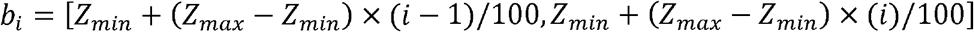; *Z_max_* is either 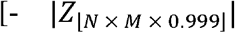 or 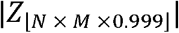 and *Z_min_* is either -2 or 2) and the percentage of readouts scored above the Z-cutoff in each bin is determined.

#### Calculation of ζ and weighted ζ score

When a screen includes a large number of both negative and positive controls, these controls are all displayed in a Zeta plot. Radial kernel SVM is next constructed to maximally separate positives from negatives in the prior defined Z range using e1071 packages of R. To avoid overfitting, it is important to use an independent dataset, such as non-expressors as internal negative controls, to confirm the deduced SVM. To provide a value to represent the regulatory function of gene i that generates a curve above the SVM curve, the area between the two curves is calculated as the Zeta score (ξ) for this gene. To highlight hits scored at higher Z score bins, the area in each bin is multiplied with the value of Z in such bin and all adjusted areas are summed to give rise to the final weighted ξ score:

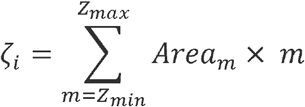

Where the *Area_m_* is the area in the specific *bin_m_*:

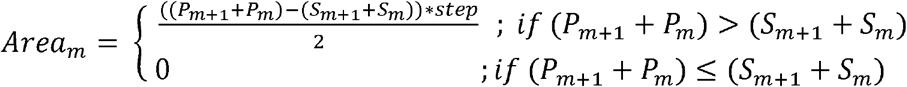

Where the *P_m_* and *P*_*m*+1_ are the y-axis values of gene i in the Zeta plot whereas *S_m_* and *S*_*m*+1_ are the y-axis values on the SVM curve, both at *bin_m_* and *bin*_*m*+1_; step is the bin size which equals to (*Z_max_* − *Z_min_*)/100.

With certain screens without any positive controls, it will be impossible to generate a SVM curve to help eliminate experimental noise. In these applications, it is still possible to calculate a ξ score for each gene by determining the *Area_m_* under the gene-specific curve at *bin_m_:*

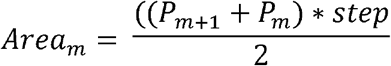

Where the *P_m,_ P_m+1_* and step are the same as those with the Area with a SVM curve.

Although ξ scores are separately generated in our application to quantify the contribution of a given gene to exon inclusion or skipping, the absolute values of these ξ scores may also be summed to reflect the global activity of such gene in regulated splicing. ZeteSuite generates this summed value as the default data output unless users select “-c no” to separately generate two ξ scores in opposition directions.

#### Screen Strength and determination of the threshold for hit selection

The ξ scores can be used to rank genes and the next important step is to define a suitable cutoff to define hits at different confidence levels. For this purpose, the concept of Screen Strength is first introduced:

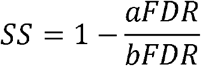

Where aFDR (apparent FDR) is the number of non-expressors identified at hits divided by the total number of hits and bFDR (baseline FDR) is the total number of non-expressors divided by all screened genes.

Next, the Screen Strength plot is generated: ξ scores are first divided into 100 even bins from the smallest to the largest and the SS value is determined at each bin. Connecting individual SS values then generates a simulated SS curve, based on which to deduce individual balance points (BPs). Users may choose one or multiple BPs to identify hits at the different SS intervals.

### ZetaSuite Part 3

#### Removing off-targeting hits

In the genome-wide screening, siRNAs were designed to specifically degrade mRNA transcripts of complementary sequence to reduce the expression of gene products. In practice, these reagents exhibit a variable degree of suppression of the targeted gene and may also suppress genes other than the intended target. The reagent’s phenotypic effects resulting from suppression unintended genes was called off-target. The reason to off-targets is due to the part-sequence complementary such as the microRNA-like off-targeting. And the consequence of off-targets is the phenotype or the effects on the readouts mainly due to off-targeting to a function gene. Multiple methods have developed to deal with the off-targeting problem based on the reason (refer DEMETER2) and consequence (refer ATARiS). Different from the many-to-one traditional screening data, the HTS^2^ can better evaluate the phenotype consistency by comparing the similarity effects on all the readouts. Based on these conditions, we defined the off-targeting hits by combine the off-targeting reason and consequence together by comparing the hits with user-defined well-known genes or total-defined hits) the off-targeting genes should have one of the targeting RNAs targeted to the well function genes (at least 11nt complementary sequence in the targeting RNA). 2) they should show high similarity on the readouts’ effects with targeted well function genes (Pearson correlation score > 0.6).

#### Functional interpretation of identified hits

Based on selected hits, ZetaSuite combine two gene function databases to interpret the functions. One is Gene Ontology database, we used ClusterProfiler to enrich hits on the GO terms. The top 15 GO terms with lowest adjust p-values were presented. The other is the CORNUM complex databases, we annotated the hits to the complexes. The top 15 complexes with highest hits’ number were gave. If the complexes number were lower than 15, the complexes with hits’ number larger than 3 would be outputted.

#### Network construction

We modified the SC3 method by using the absolute values of Spearman and Pearson correlation score to calculate the distance matrix and then used it to do the clustering. After SC3 analysis, each gene pair has a consensus score, which measures the regulation strengths. The consensus scores were then used as the edge weights. Gene-gene similar correlation and anti-correlation were annotated as different edge types. The nodes in the network represent the hits identified by ZetaSuite pipeline. Larger nodes’ size means larger ζ value. The color of the nodes act for the clusters based the SC3 calculation. Cluster number was chosen based on the total within-cluster sum of square “elbow” site. The resultant hit networks were visualized with Gephi by using a Yifan Hu Proportional layout. Disconnected nodes were then trimmed from the graph before generating the plots.

### Other experimental procedures

#### Testing the multiple testing correction methods on error rate reduction

The multiple testing correction methods, as FDR, Bonferroni correction etc., are frequently used to reduce error accumulation in multiple hypothesis testing. However, it can only be used to deal with the data from one-dimensional screens, but not suitable for two or multiple dimensional screens. To further test this, a common cutoff is Z>=3 or <= –3, and thus, the estimated false positive level (p-value) is below 0.01, meaning that for each readout, a given siRNA has 1% chance to be identified as a false positive hit. For all conditions, we did ∼15,000 tests for each readout, and using the most stringent Bonferroni correction, we obtained a corrected p-value of 0.01/15000=6.67×10^-7^ and a corresponding Z=4.97. Now using Z=4.97 as the corrected cutoff to choose hits, we found that the false positive level was still as high as 24.9%. We therefore concluded that such canonical multiple testing correction methods are not sufficient to reduce the accumulation of errors with increasing readouts in two-dimensional high throughput screens.

#### Evaluating the optional number of functional readouts in two-dimensional screen

Positive controls and high-confidence hits, the latter of which are defined based on total readouts, are used as references in our evaluation. The number of readouts is progressively down-samples to 50, 100, 150, 200, 250 and 300 using R Sample function without replacement and each specific number of down-sampled readouts are replicated 3 times. Down-sampled matrixes are processed using the same ZetaSuite pipeline. Hits from down-sampled matrixes are used to determine the percentage of the hits over the reference sets.

#### Analysis of the splicing screen data with RIGER

RIGER was originally developed to identify essential genes in genome-scale shRNA screens ^17^. In RIGER, the signal-to-noise ratio is entered as input, which is now replaced with the Z-scores for individual alternative splicing readouts. The data are then processed with the latest version of RIGER (2.0.2) from the website as provided in the source table above. Default RIGER parameters are used in all steps, except that the number of permutations is set to 100,000 to obtain a more precise *p*-value for each pool of siRNAs. The FDR is computed from the empirical permutation p-values using the Benjamini-Hochberg procedure. This enables ranking of siRNA pools by FDR.

#### Analysis of splicing screen data with RSA

RSA is a probability-based method to identify hits, requiring data generated with multiple targeting siRNAs against each gene ^16^. In RSA, fold-changes of treated over control samples are entered as input. In our application, the inputs are fold-changes of the splicing ratio of a given alternative splicing event in a siRNA pool-treated well divided by the averaged splicing ratio from NS-mix treated wells. The entered data are processed with the latest RSA software, as specified in the source table above. The following parameters -l 0.2 -u 0.8 and -l 1.2 -u 2.0 are used to select hits for induced exon inclusion and skipping, respectively.

#### Analysis of splicing screen data with MAGeCK

MAGeCK is a statistical method designed to quantify the collective activity of multiple siRNAs against each gene by using the robust rank aggregation (RRA) algorithm ^18^. In order to meet the MAGeCK input requirement, each Z-score in the ZetaSuite input matrix is first converted to p-value. The input data are processed with the modified RRA algorithm, as in MAGeCK, with default parameters.

#### Processing DRIVE and DepMap cancer dependency datasets

The DRIVE and DepMap data already processed with DEMETER2 are downloaded from https://depmap.org/portal/download/. DepMap generated 3 independent datasets. In order to avoid experimental variations in different datasets, only the biggest DepMap dataset is selected for current analysis, which includes 285 cancer cell lines across approximately 100k shRNAs. ZetaSuite is applied to this dataset to calculate weighted ζ-scores with the parameters -z no –svm no and -c no.

#### Feature association analysis on cancer dependencies and checkpoints

To analysis association with CNV or gene expression, cancer cell lines are ranked based on the levels of CNV in a given gene or expression of the gene. Cancer dependency scores are next compared between cell lines in top 25% versus bottom 25% and Wilcox-test is performed to determine the *p*-value for the gene. To analysis association with mutations, cancer cell lines are divided in two groups with or without mutation in each gene. The cancer dependency scores are next compared between these two groups and Wilcox-test is performed to generate the *p*-value for the gene.

### Data and code availability

The datasets used to evaluate the existing and new designed methods are available at the website:XXX. The open source ZetaSuite is freely available at website XXX. We will update this website periodically with new versions.

## Acknowledgements

This work was supported by grants from NIH grants HG004659 and DK098808. We also wish to thank multiple former trainees in the Fu lab, particularly, Drs. Yu Zhou, Hairi Li, Jinsong Qiu, Bing Zhou, and Xuan Zhang for their contributions to the ideas that led to the development of the ζ statistic. We are also grateful to Dr. Shirley Liu of Dana Farber Cancer Institute for critical reading and comments on the manuscript.

## Contributions

Y.H. was responsible for all bioinformatic analysis and software development. C.S. contributed to data interpretation and presentation and G.Z. provided various assistance in developing software. Y.H., and X.F. wrote the manuscript. All authors edited the paper.

## Supplementary information

Supplementary Figs. 1-6

