## Supplementary materials for "ZetaSuite, A Computational Method for Analyzing Multi-dimensional High-throughput Data, Reveals Genes with Opposite Roles in Cancer Dependency"

### Supplementary Figure 1

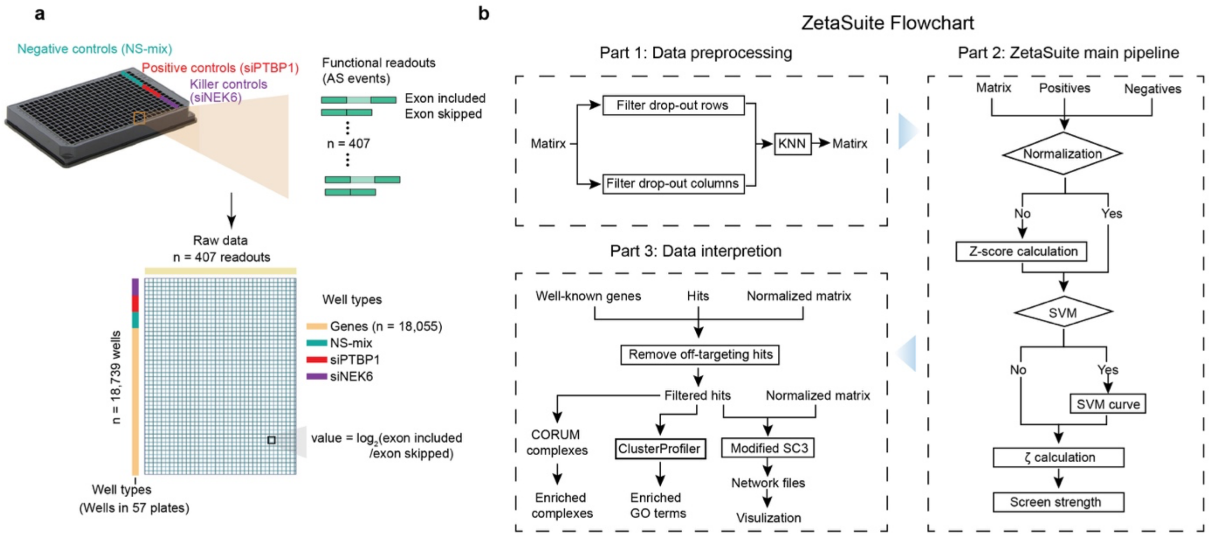


**Supplementary Figure 1. Overview of in-house data set and the ZetaSuite flowchart.**

**a**, In-house data format. Two-dimensional in-house dataset was generated from a siRNA screen to identify splicing regulators. In each siRNA-treated well, 407 pairs of alternative splicing (AS) events are interrogated by RNA Annealing Selection Ligation sequencing (RASL-seq), as detailed elsewhere. A total number of 18,480 siRNA pools against annotated protein-coding genes are arrayed in 57 384-well plates. Each plate contains 6 negative controls (NS-mix), 5 positive controls (siPTBP1) and 5 killer controls (siNEK6). After screening, the raw data are tabulated in a matrix as the $log_2$ isoform ratio (exon included isoform/exon skipped isoform). **b**, The flowchart of the ZetaSuite software in three parts (https://github.com/YajingHao/ZetaSuite), as detailed in the text.

### Supplementary Figure 2

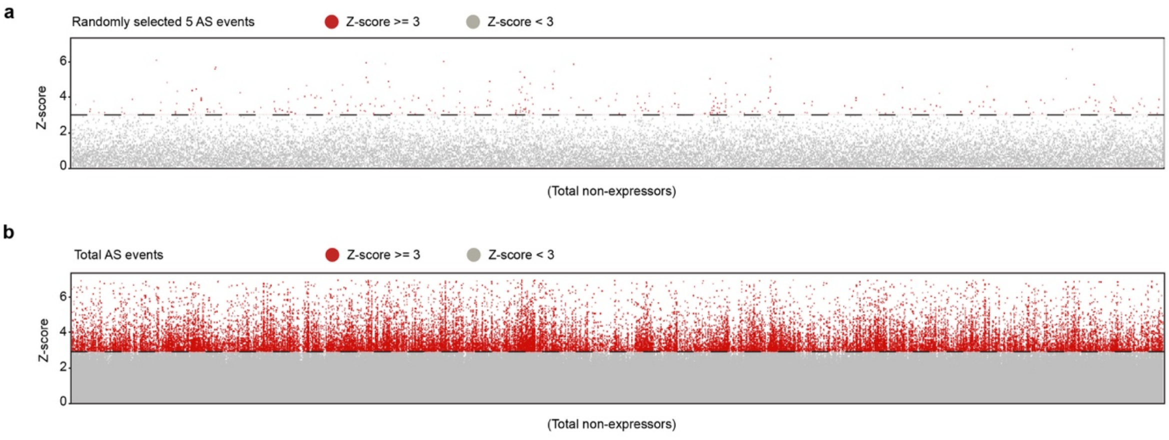


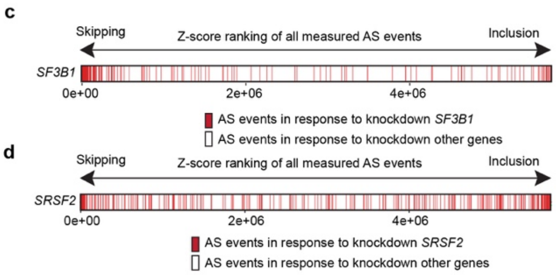


e

| Methods | | FDR cutoff | | | |
|---|---|---|---|---|---|
| | | 1% | 5% | 10% | 25% |
| RSA | Skipping | 3,436 | 4,494 | 5,136 | 6,367 |
| | Inclusion | 3,482 | 4,781 | 5,781 | 7,572 |
| RIGER | Skipping | 826 | 1,225 | 1,473 | 2,012 |
| | Inclusion | 1,802 | 2,373 | 2,727 | 3,393 |
| MAGeCK | Skipping | 2,231 | 2,608 | 2,819 | 3,205 |
| | Inclusion | 2,105 | 2,560 | 2,805 | 3,207 |

**Supplementary Figure 2. Data analysis using existing statistical approaches.**

**a**-**b**, The distribution of Z-scores of all non-expressors (n=5006) based on 5 randomly selected AS events (**a**) or all interrogated AS events (**b**). Red-marked dots indicate hits with Z>=3, showing the majority (~80%, see Fig. 2**c**) of non-expressors scored as false-positive hits when all measured AS events were included in the analysis with the traditional Z-based approach. **c**-**d**, The Z-score rank distribution (from induced exon skipping on left to exon inclusion on right) of *SF3B1* responsive AS events (**c**) and *SRSF2* responsive AS events (**d**) among total detected AS events in the screen, showing skewing of *SF3B1*-induced splicing toward exon skipping and *SRSF2*-induced splicing in both directions. **e**, The hits number at some common FDR cutoffs using 3 different existing methods.

**Supplementary Figure 3**

**a**

| Gene symbol | PMID | Gene function |
|---|---|---|
| SRSF3 | 31462429 | serine and arginine rich splicing factor 3 |
| SRSF2 | 23562324 | serine and arginine rich splicing factor 2 |
| HNRNPK | 22684629/11867641 | heterogeneous nuclear ribonucleoprotein K |
| HNRNPL | 19946215 | heterogeneous nuclear ribonucleoprotein L |
| SRSF7 | 25482508/17137791 | serine and arginine rich splicing factor 7 |
| FUS | 22829983/24204307 | FUS RNA binding protein |
| SON | 21504830/20705336 | SON DNA binding protein |
| EIF4A3 | 25476502 | eukaryotic translation initiation factor 4A3 |
| U2AF2 | 25326705 | U2 small nuclear RNA auxiliary factor 2 |
| YJU2 | 28373290 | YJU2 splicing factor homolog |

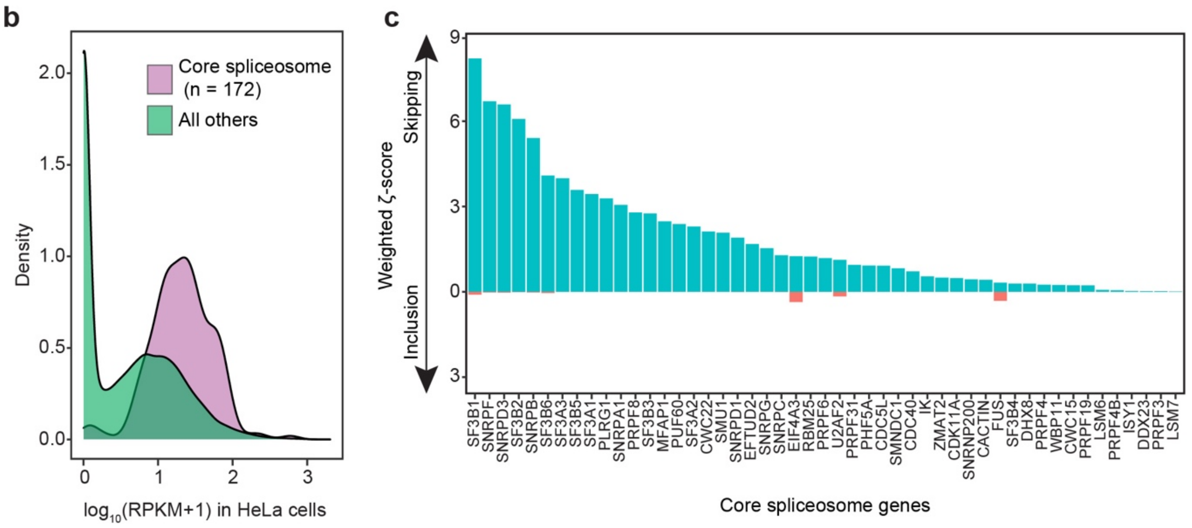


**Supplementary Figure 3. Use of weighted ζ-scores to characterize screen hits.**

**a**, The list of 10 known splicing regulators displayed in the Zeta plot in Fig. 3**a**,**b**, The density of gene expression levels for annotated core spliceosome genes compared to all other genes in HeLa cells. **c**, Weighted ζ-scores of representative core spliceosome genes in induced exon skipping (blue) or inclusion (red), showing that knockdown of core spliceosome components predominately induced exon skipping.

### Supplementary Figure 4

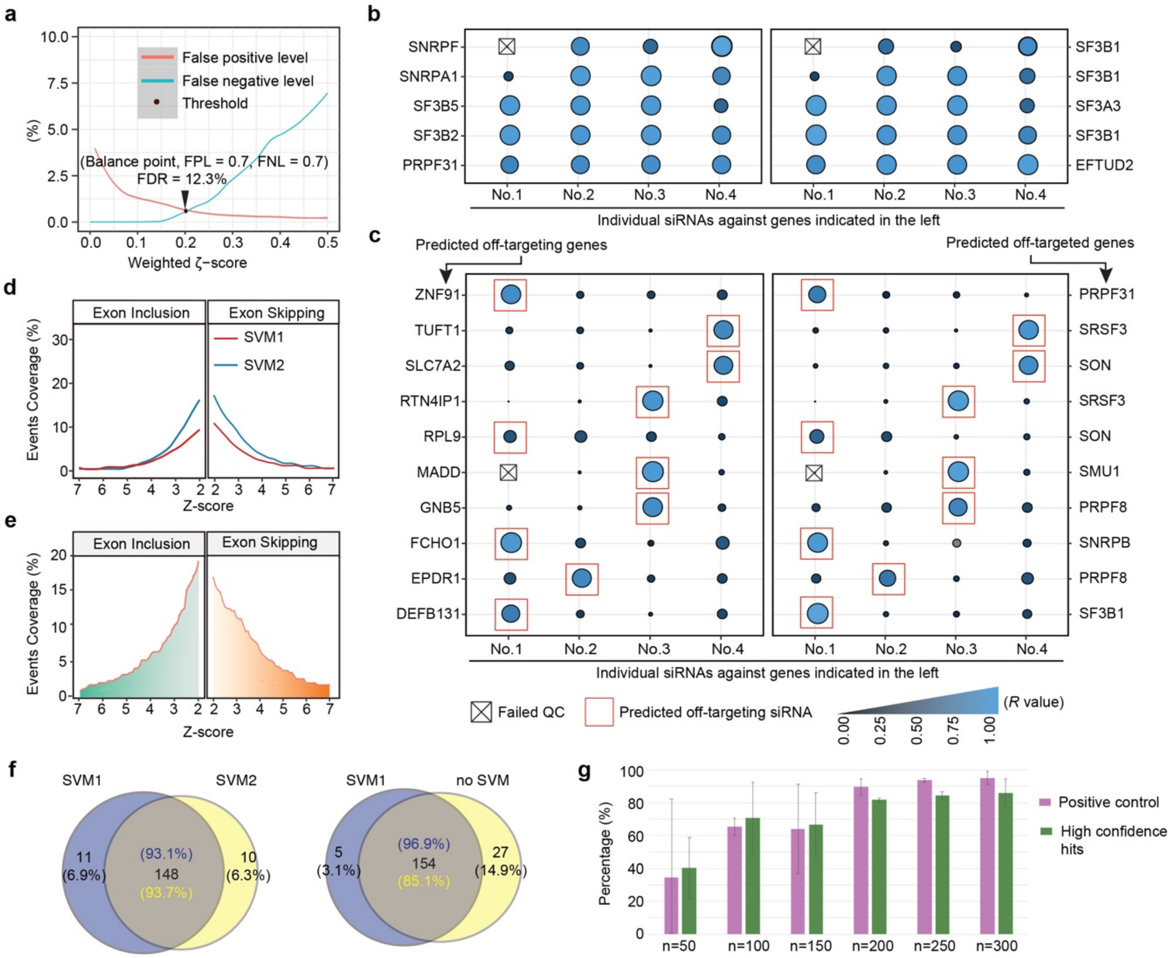


**Supplementary Figure 4. Strategy to remove off-target effects and optimal readouts for two-dimensional genome-wide screens.**

**a**, The balanced error level (BRL) approach for setting the threshold. Arrow indicates the chosen threshold and associated FDR. **b**-**c**, Hits with related functions or due to off-target effects. Results of secondary screen with 4 individual siRNAs in comparison with the pool of those siRNAs (left) or with the pools of other siRNAs against genes with related functions (right). The size of circles reflects the level of similarity in splicing response. Hits are due to related functions when multiple single siRNAs produce similar results (**b**) or to off-target effects when a single siRNA is responsible for the similarity to both siRNA pools (**c**). **d**, Deduced SVM curves using two different sets of positive controls. SVM1 was defined with siPTBP1 repeats and SVM2 with a set of known spliceosome components listed in Supplementary Fig. 3**a**. **e**, Diagram to illustrate the calculation of weighed ζ-scores without using a SVM curve. **f**, Venn diagrams showing the overlaps of high confidence hits selected using different SVM curves (left) or with and without using a SVM curve (right). **g**, The impact of readout (AS event) size on the efficiency in recovering a set of reference hits. Each bar represents the percentage of recovered reference hits (purple for siPTBP1 replicates; green for high confidence hits based on total AS events) by ZetaSuite using different numbers of readouts. Error bars represent the standard deviation from three independent samplings.

**Supplementary Figure 5**

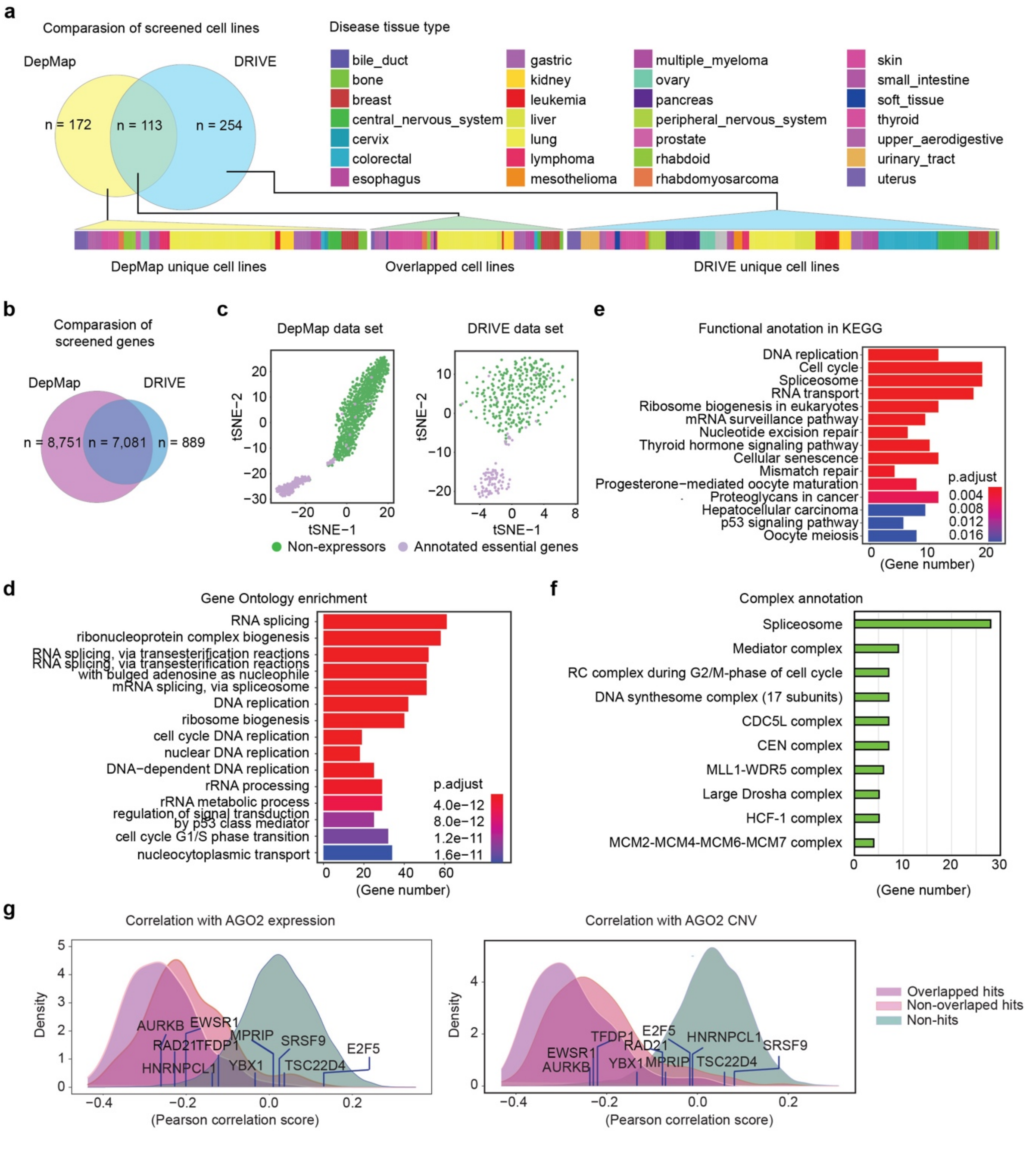


**Supplementary Figure 5. Significantly increased number of fitness genes identified by ZetaSuite from the existing DepMap and DRIVE datasets.**

**a**, Comparison of cell lines surveyed by DepMap and DRIVE projects. Cell lines derived from different cancer types are color-indicated. A common set of 113 cell lines was analyzed by both projects. **b**, Comparison of genes interrogated by the two projects. **c**, Robust segregation of positive (purple: annotated essential genes) and negative (green: non-expressors) controls in DepMap (left) and DRIVE (right). **d**, GO term enrichment for newly identified essential genes by ZetaSuite. **e**, The function enrichment on KEGG pathways for newly identified essential genes by ZetaSuite. **f**, Top 10 enriched complexes of newly identified essential genes by ZetaSuite. Complexes are from the CORUM database. **g**, The density plots of correlation between DEMETER cancer dependency score and AGO2 expression (left) or copy number variation(right) for different gene sets according to the color key on right. The overlapped and non-overlapped hits correspond to those displayed in Fig. 5**e**. Ten genes uniquely detected by DRIVE are labeled, showing that 8 of 10 are distributed with non-hits.

### Supplementary Figure 6

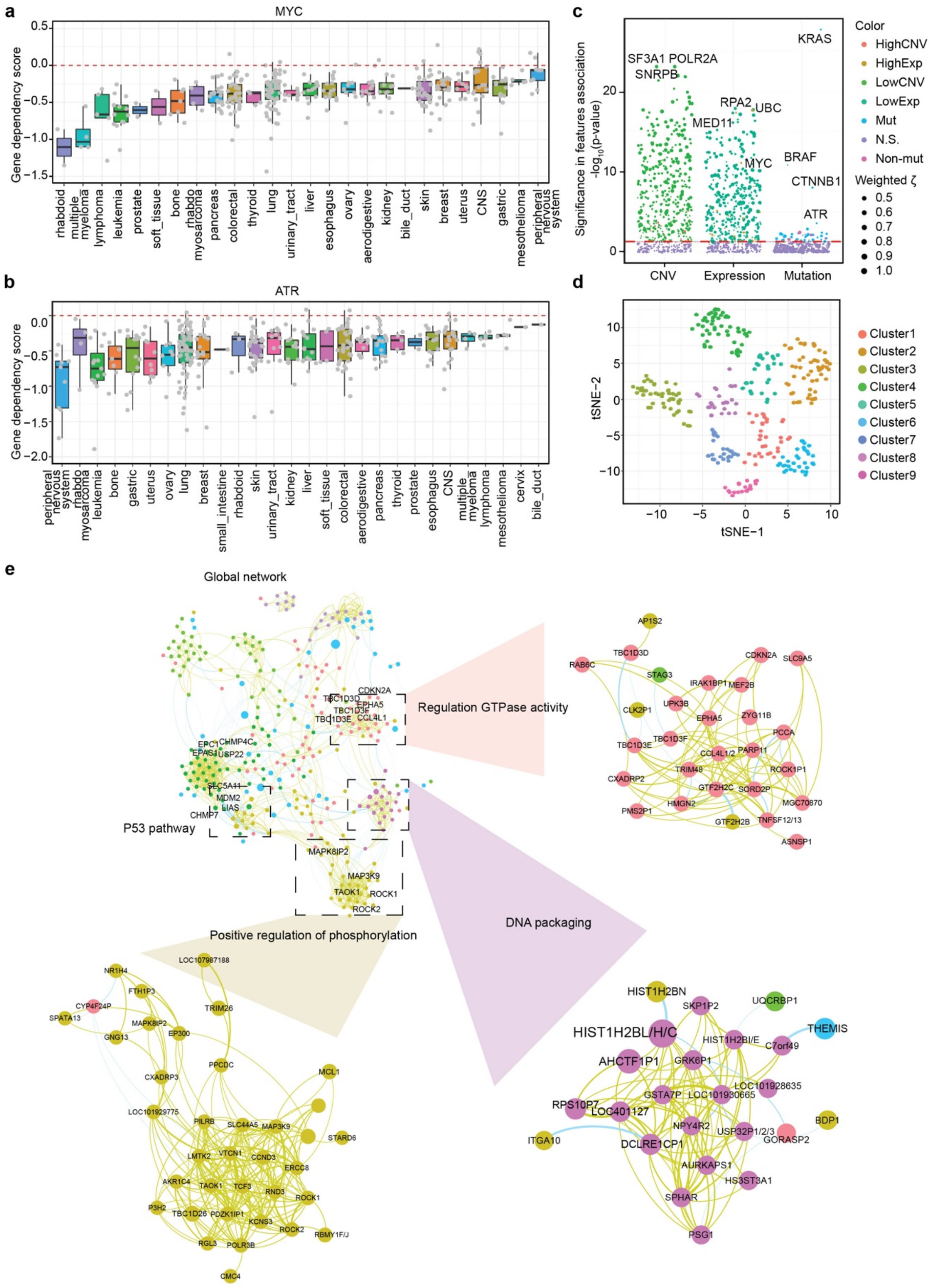


**Supplementary Figure 6. Functional analysis of identified hits by ZetaSuite.**

**a**-**b**, Averaged dependency scores of *MYC* (**a**) and *ATR* (**b**) in different cancer tissues. **c**, The association of ZetaSuite-identified cancer dependencies with gene expression, copy number and mutation features as in Fig. 6**e**. **d**, Clusters of hits detected by ZetaSuite that led to improved tumor cell proliferation. **e**, Global network of tumor checkpoint hits. Highlighted sub-networks include those involved in the regulation of GTPase activities, DNA packaging, and protein phosphorylation.
